# Scalable proteomics with multiDIA resolves proteoforms in neurodegenerative disease

**DOI:** 10.64898/2026.10.16.756994

**Authors:** Orhun Kok, Corinna Friedrich, Kirsten Altig, Derek Oakley, Bradley T. Hyman, Mahlon Collins, Nikolai Slavov

## Abstract

Deep proteome characterization has traditionally required multidimensional chromatographic fractionation, long acquisition times and large sample amounts, thus limiting scalability. Here, we introduce multiDIA, which combines independent digestions with multiple proteases and data-independent acquisition (DIA) to increase sequence coverage without extensive chromatographic fractionation. By generating complementary and overlapping peptide representations of proteins, multiDIA identifies about 270,000 distinct peptides per sample, including 3,649 phosphosites. This is more than twice the coverage of unfractionated workflows and exceeds the coverage of multidimensional fractionation workflows that use 10-fold more material and time. We demonstrate the approach in Parkinson’s disease (PD) brain samples, showing that multiDIA substantially expands sequence coverage while retaining the sensitivity, reproducibility, and scalability of DIA. This enabled the identification of disease-associated proteoform changes undetectable by single-protease DIA and the quantification of many disease-relevant post-translational modifications. MultiDIA thus enables deep, high-confidence proteome characterization from limited sample quantities without the throughput and sensitivity constraints of multidimensional fractionation.

## Introduction

Mass spectrometry–based proteomics encompasses a diverse range of workflows that trade proteome depth and sequence information against analytical throughput. At one extreme, comprehensive proteome characterization has traditionally relied on extensive multidimensional chromatographic separation, often including fractionation followed by separate analysis of each fraction^1–3^. These approaches can achieve deep sequence coverage but substantially increase analysis time. They typically use data-dependent acquisition (DDA), which selects and fragments individual precursors and thus generates relatively simple spectra that support confident identification of amino acid sequences and post-translational modifications^4,5^. However, the serial nature of precursor selection and the need to analyze many fractions make such workflows difficult to scale to large numbers of samples.

Recent advances in sample preparation, data acquisition and interpretation^6–8^, particularly for data-independent acquisition (DIA), have substantially improved proteomics sensitivity and scalability^9,10^. By systematically fragmenting broad precursor populations rather than selecting individual precursors, DIA supports reproducible analysis of many samples without the extensive fractionation required by workflows using multidimensional chromatography for deep sequence coverage. Yet this scalability comes with an important limitation: DIA workflows without extensive chromatographic separation generally provide less comprehensive sequence coverage (about 120,000 peptides per sample) than deeply fractionated workflows (over 225,000 peptides per sample)^11^. Furthermore, the multiplexed nature of DIA spectra can make confident assignment of peptide fragments – and thus sequences and modifications – more challenging. Thus, a fundamental trade-off remains between scalable proteome measurement and comprehensive, high-confidence sequence characterization. Much higher sequence depth has been achieved by combining six different proteases, multidimensional chromatographic separation and different peptide fragmentation methods, but the long analysis times and large sample requirements have limited this approach to a few cell lines^3^.

Here, we sought to mitigate the trade-off between throughput, identification confidence and sequence coverage by combining the sensitivity and scalability of DIA with the complementary sequence information provided by independent digestion with multiple proteases in a workflow we term multiDIA. Rather than increasing proteome depth through extensive chromatographic fractionation, we generate orthogonal peptide representations of each sample by digesting aliquots independently with different proteases and analyzing each digest by DIA. While tryptic peptides generally yield the deepest protein sequence coverage, this coverage remains incomplete because many protein regions, such as RS domains and low-complexity domains, do not produce detectable tryptic peptides^12^. To complement trypsin and increase protein sequence coverage, multiple proteases have long been used^13–17^ but have not yet been combined with modern DIA to establish the feasibility of deep and scalable proteomics workflows.

Multiple protein isoforms, termed proteoforms, often emerge from the same gene, including splice isoforms, translationally, and post-translationally modified forms^18,19^. These proteoforms often have distinct biological functions but may not be distinguishable by MS analysis of tryptic digests, owing to their limited sequence coverage^20^. Moreover, quantification that summarizes all peptides of a gene into a single abundance averages over its proteoforms and can mask changes specific to one of them^21^. As a result of these limitations the role of specific proteoforms in human health and disease remains poorly understood.

To benchmark the ability of combining multiple proteases with DIA to identify disease-relevant proteoforms, we profiled bulk brain tissue samples from a cohort of Parkinson’s disease (PD) and healthy control samples. Multiple proteoforms are implicated in PD pathogenesis. For example, alpha-synuclein splice isoforms and PTMs alter the protein’s aggregation propensity thereby contributing to the emergence of the disease’s hallmark Lewy body pathology. Tau co-pathology also frequently occurs in PD, but the specific tau proteoforms most strongly associated with PD remain uncharacterized. α-synuclein (SNCA) has splice isoforms lacking exon 3 or exon 5^22^, and the isoforms of tau (MAPT), which differ by the inclusion of exons are all expressed in the adult human brain^23,24^. More generally, the brain exhibits particularly high levels of alternative mRNA splicing^25,26^, which generates additional proteoform diversity. PD is associated with widespread proteomic changes, including those resulting from loss of protein homeostasis, inflammation, activation of cell death pathways, and axonopathy^27^. Yet standard workflows combining two-dimensional chromatography with tandem mass tags (TMT) have quantified only 50,000–80,000 unique peptides per PD brain sample, limiting proteoform resolution^28^.

Another key motivation for multiDIA is to increase confidence in assigning amino acid sequences and modifications through redundant sequence coverage. Digesting proteins with different proteases produces overlapping peptides representing the same protein regions. This redundant representation of protein sequences by overlapping peptides can provide orthogonal evidence for the corresponding sequence assignments, increasing confidence in sequence identification and in the characterization of sequence variants and post-translational modifications. This increased confidence can support the confident identification of modified peptides and the discovery of new proteoforms, such as those arising from alternate RNA decoding^29^.

We tested this strategy on dorsolateral striatum samples from a cohort of PD and control cases, evaluating whether multiDIA can approach the depth and sequence confidence traditionally achieved with extensively fractionated samples while retaining the throughput and scalability of DIA. Using about 3 hours of analysis time per sample, we identified about 270,000 peptides per brain sample, exceeding the coverage of a previous workflow that identified up to 195,000 peptides per TMT set by analyzing 120 offline fractions, each on a 2- or 3-hour online gradient (about 300 hours of analysis time per TMT set)^30^. This increased sequence coverage allowed us to detect more proteoforms and phosphosites than tryptic peptides alone, including those of PD-associated proteins alpha-synuclein and dopa decarboxylase.

## Results

### MultiDIA increases proteome sequence coverage

To characterize proteome changes in PD, we analyzed bulk dorsolateral striatum tissue from 12 donors (6 PD cases and 6 controls; Table 1). Samples were dissected and prepared using single-pot solid-phase-enhanced sample preparation (SP3)^31^. Each sample lysate was split into three aliquots, which were digested separately with trypsin, Lys-C, or Glu-C to generate complementary peptide populations and increase proteome coverage (Fig. 1a). The resulting peptides from each digest were analyzed by DIA using a 60-window dia-PASEF method on a timsTOF Ultra mass spectrometer. One Lys-C sample was lost during handling and was not acquired, and two Lys-C samples, one Glu-C sample and one trypsin sample had abnormally low coverage owing to partial sample loss.

**Table 1.**
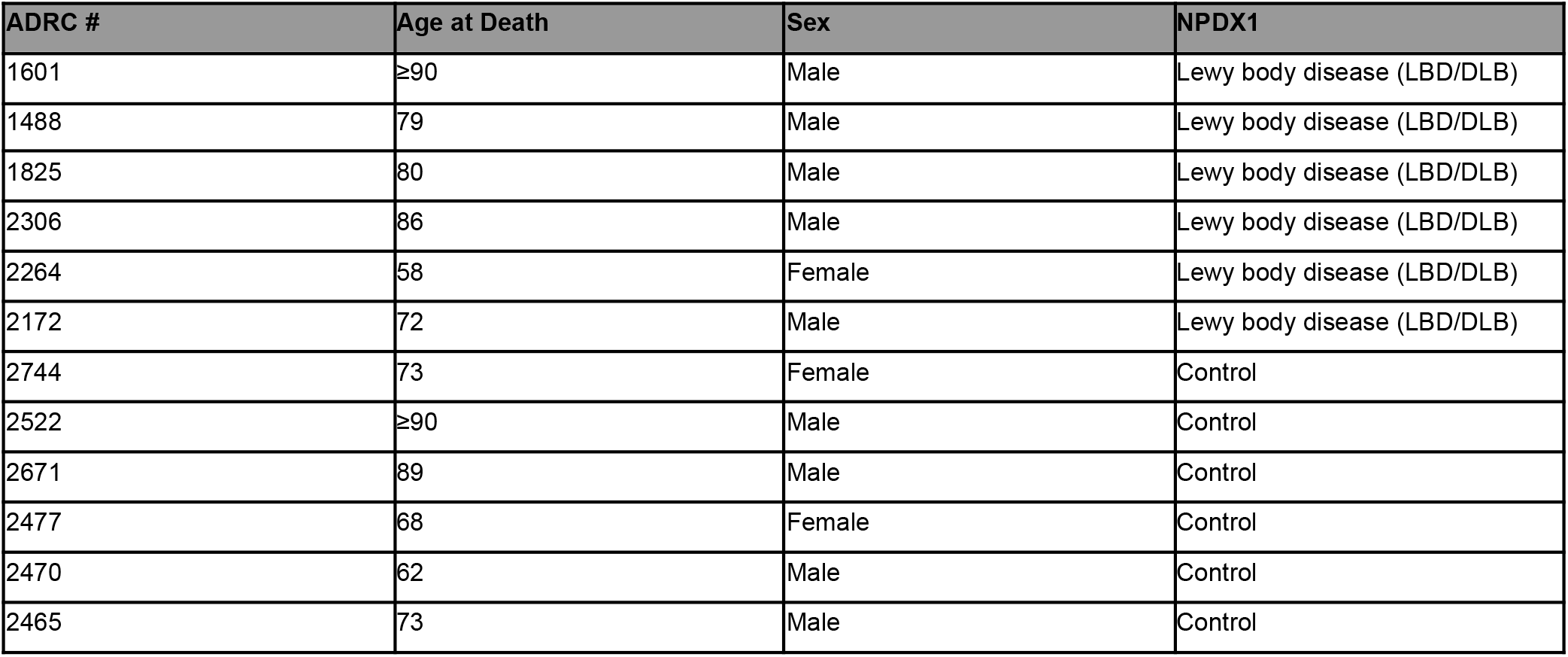
Metadata for the PD and control donors used in this study. All samples came from the Massachusetts Alzheimer’s Disease Research Center (ADRC) brain bank.

| ADRC # | Age at Death | Sex | NPDX1 |
| --- | --- | --- | --- |
| 1601 | ≥90 | Male | Lewy body disease (LBD/DLB) |
| 1488 | 79 | Male | Lewy body disease (LBD/DLB) |
| 1825 | 80 | Male | Lewy body disease (LBD/DLB) |
| 2306 | 86 | Male | Lewy body disease (LBD/DLB) |
| 2264 | 58 | Female | Lewy body disease (LBD/DLB) |
| 2172 | 72 | Male | Lewy body disease (LBD/DLB) |
| 2744 | 73 | Female | Control |
| 2522 | ≥90 | Male | Control |
| 2671 | 89 | Male | Control |
| 2477 | 68 | Female | Control |
| 2470 | 62 | Male | Control |
| 2465 | 73 | Male | Control |

**Figure 1.**
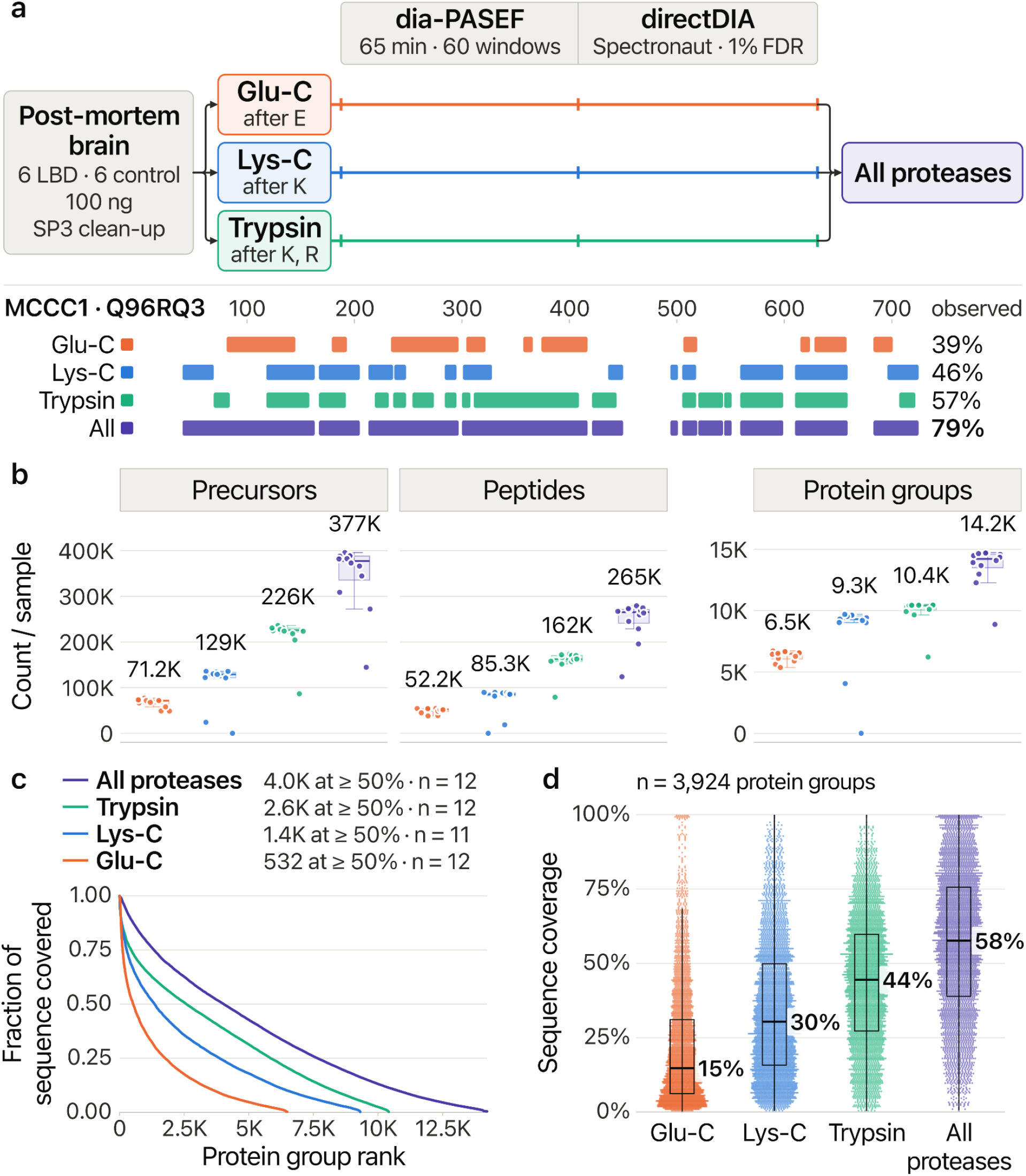
multiDIA workflow and sequence coverage. (a) Workflow schematic: aliquots of each sample are digested separately with Glu-C, Lys-C or trypsin; each digest is analyzed and searched separately, and the results are pooled (top). Below, the observed coverage of MCCC1 across all runs: trypsin alone covers 57% of the sequence and the three digests together cover 79%, with each digest adding residues missed by the others. **(b)** Number of quantified precursors, peptides and protein groups per sample. **(c)** Protein groups ranked by sequence coverage: 4.0K reach ≥50% coverage with all digests pooled, versus 2.6K with trypsin alone. **(d)** Sequence coverage of the 3,924 protein groups detected in all three digests; the same proteins are covered more deeply by multiDIA.

Relative to trypsin alone, multiDIA increased the number of precursors identified per sample 1.7-fold (median 377,410 versus 225,996) and the number of distinct peptides 1.6-fold (265,366 versus 162,238; Fig. 1b). The larger number of quantified peptides per sample also increased the number of quantified proteoform groups, from about 10,400 with trypsin alone to more than 14,000 (Fig. 1b). The number of protein groups with at least half of their sequence covered increased 1.6-fold, from 2,602 to 4,043 (Fig. 1c). For the 3,924 protein groups detected by all three proteases, median sequence coverage rose from 44.5% with trypsin alone to 57.6% with multiDIA (Fig. 1d).

## Increased sequence coverage resolves proteoforms

Each search assigns a peptide to a protein group, which can include one or more proteoforms; thus, the protein group alone does not reveal which proteoforms a peptide derives from. We therefore mapped each peptide to proteoforms by searching for its exact sequence in the canonical (as defined by UniProt) form and in every annotated proteoform originating from the corresponding gene, noting which digest had identified it. Aligning each proteoform to its canonical form defined its discriminating region (the residues absent from the canonical form) and its deletion junctions (the points where canonical residues are missing). Only peptides absent from the canonical sequence were counted as proteoform evidence.

Each protease identified sequences that helped resolve proteoforms, and combining the proteases increased the coverage of proteoform-specific sequences. For each non-canonical proteoform, we measured how much of its discriminating region (residues absent from the canonical sequence) the peptides covered, and whether a peptide spanned a splice junction (the position at which a segment of the canonical sequence is spliced out) by covering residues on both sides of it. Proteoforms that differ from their canonical form only by a shorter N or C terminus have neither feature and were not counted. Across the 1,307 non-canonical proteoforms with at least one discriminating residue covered, multiDIA raised the median coverage of the discriminating region to 62% compared to 38% with trypsin alone (Fig. 2a). It also increased 1.4-fold the number of proteoforms with at least half of their discriminating region covered (762 versus 545) and 1.3-fold the number with at least one junction-spanning peptide (757 versus 584; Fig. 2a). Here we note that covering most of a discriminating region is harder than spanning a deletion junction: discriminating regions are often longer than a single peptide, whereas one peptide can span a junction.

**Figure 2.**
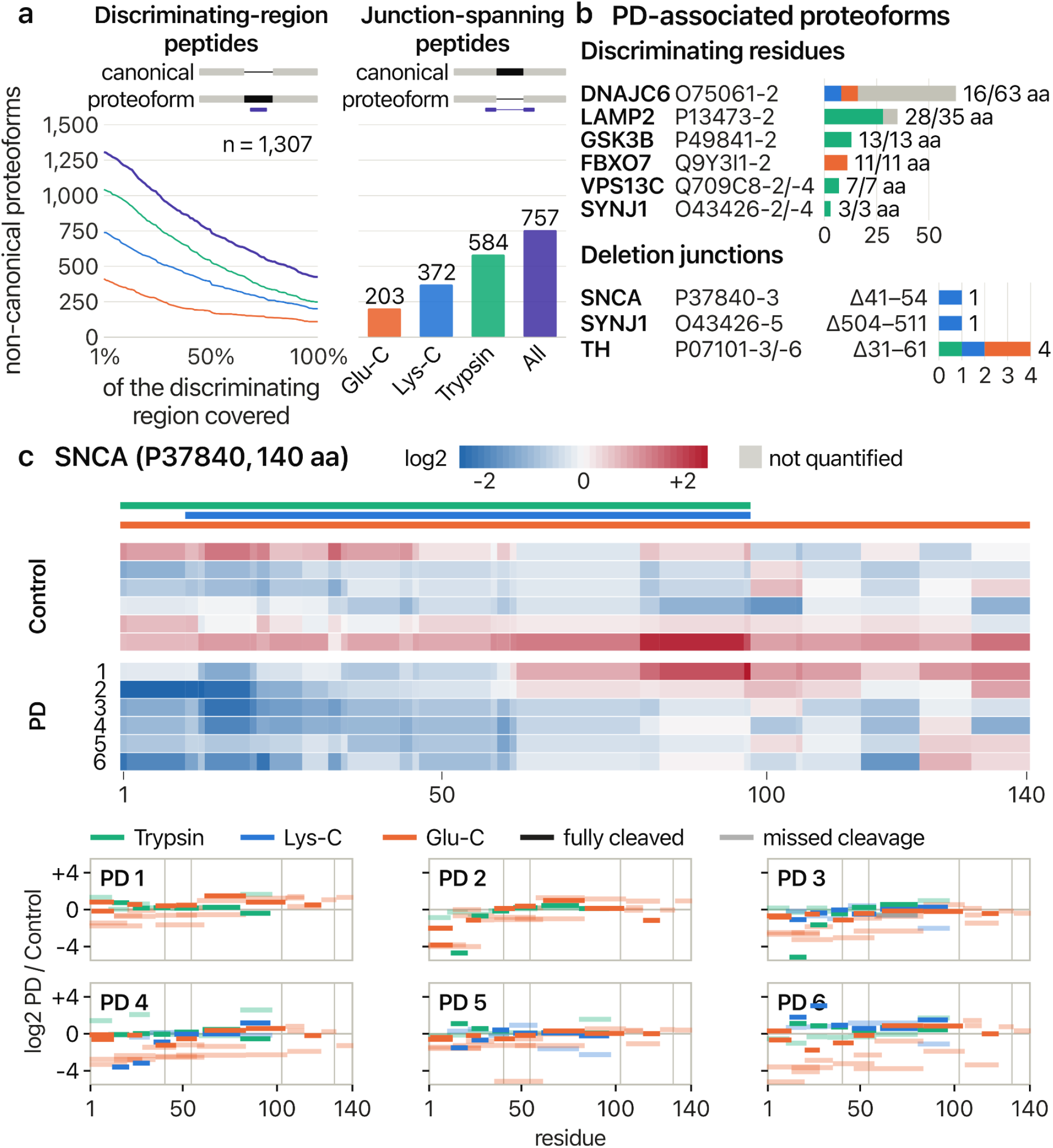
Proteoforms and PD-associated peptides by donor. (a) Proteoforms by discriminating-region coverage and with a junction-spanning peptide. **(b)** Proteoforms of PD-associated genes; grey, not covered. a and b count only peptides absent from the canonical sequence. **(c)** α-Synuclein (SNCA) peptide abundance by donor, log2 relative to the control mean (each control against the other controls). Top: one row per donor, ordered within each group by profile similarity; each residue shows the mean of its covering peptides; grey, not quantified. Bottom: each PD donor’s peptides as lines over the residues they cover; vertical lines mark the deletions of P37840-3 (Δ41–54) and P37840-2 (Δ103–130).

Among proteoforms of PD-associated genes, Glu-C and Lys-C peptides covered discriminating residues or deletion junctions of DNAJC6, FBXO7, SNCA and SYNJ1 proteoforms that were missed by trypsin, and peptides from all three proteases spanned the deletion junction of the TH proteoforms P07101-3/-6 (Fig. 2b). For tau (MAPT), junction-spanning peptides from all three proteases independently identified the 0N proteoforms, in which exon 1 joins exon 4 (five peptides), and the 1N proteoforms, in which exon 2 joins exon 4 (three peptides); both junctions were detected in all 12 donors (Supplementary Fig. 3). Peptides found only in the exon 4-containing big tau proteoform, from exons 4a and 6, showed no consistent difference between PD and control donors (median log2 PD/control +0.1 to +0.3 per protease; Supplementary Fig. 3).

We sought to use our deep sequence coverage to explore truncation products of α-synuclein (SNCA). It is the main constituent of Lewy bodies, and its proteolytically truncated proteoforms have been associated with aggregation, but their direct detection is challenging^32^. To this end, we plotted the relative abundance of each peptide in each donor at the residues the peptide covers, so that abundance can be compared along the protein sequence and between donors (Fig. 2c). Glu-C peptides covered all 140 residues of α-synuclein, including the 45 residues of its acidic C-terminal tail, of which tryptic and Lys-C peptides each covered only 2; together, the three proteases quantified α-synuclein across its full length in all 12 donors. Overall, α-synuclein peptide abundance ordered donors within each group by the similarity of their peptide profiles revealed region-specific patterns: peptides from the N-terminal region (residues 1–60) were on average 2-fold below the control mean in three PD donors, whereas controls ranged from ™0.5 to +1.1 log2 (Fig. 2c, top). Plotting each PD donor’s peptides separately showed that this difference was reflected the Glu-C peptides (™1.5 log2, PD versus control), whereas tryptic and Lys-C peptides covering the same residues were close to the control mean (™0.1 log2; Fig. 2c, bottom). In some patients, such as PD2, all N terminal peptides are less abundant than in the control cases, consistent with N terminal truncation.

Despite the small cohort size, the magnitude of some differences reach robust statistical significance. For example, peptides of the dopaminergic enzymes DDC and tyrosine hydroxylase (TH) were markedly lower in four of the six PD donors, 2.0–3.5 log2 (DDC) and 2.5–3.2 log2 (TH) below the control mean, whereas controls ranged from ™0.5 to +0.5 (Supplementary Fig. 2). Testing each proteoform for proteoform-specific differential abundance, defined as its log2 PD/control ratio minus that of its canonical form, four of 1,217 proteoforms differed from their canonical form at q ≤ 0.05 (Supplementary Fig. 1): P27816-3 (MAP4), Q9ULH7-4/-5 (MRTFB), Q32MZ4-4 (LRRFIP1) and P11277-2 (SPTB).

### MultiDIA expands the detection and quantification of phosphosites

All digests were searched with phosphorylation as a variable modification. Across the 10 donors measured by all three digests, multiDIA detected 3,649 phosphosites (localization probability ≥ 0.75; median 2,127 per donor, range 1,451–2,267), 1.6-fold more than trypsin alone (2,267; median 1,288 per donor; Fig. 3a). Lys-C detected 1,698 phosphosites and Glu-C 431. The digests overlapped little: only 94 phosphosites were detected by all three, while 1,646 were detected only by trypsin, 1,087 only by Lys-C and 263 only by Glu-C (Fig. 3a). Thus, 1,382 of the 3,649 phosphosites (38%) were missed by trypsin and detected only in the Lys-C or Glu-C digests (1,119 and 263 phosphosites, respectively; Fig. 3b).

**Figure 3.**
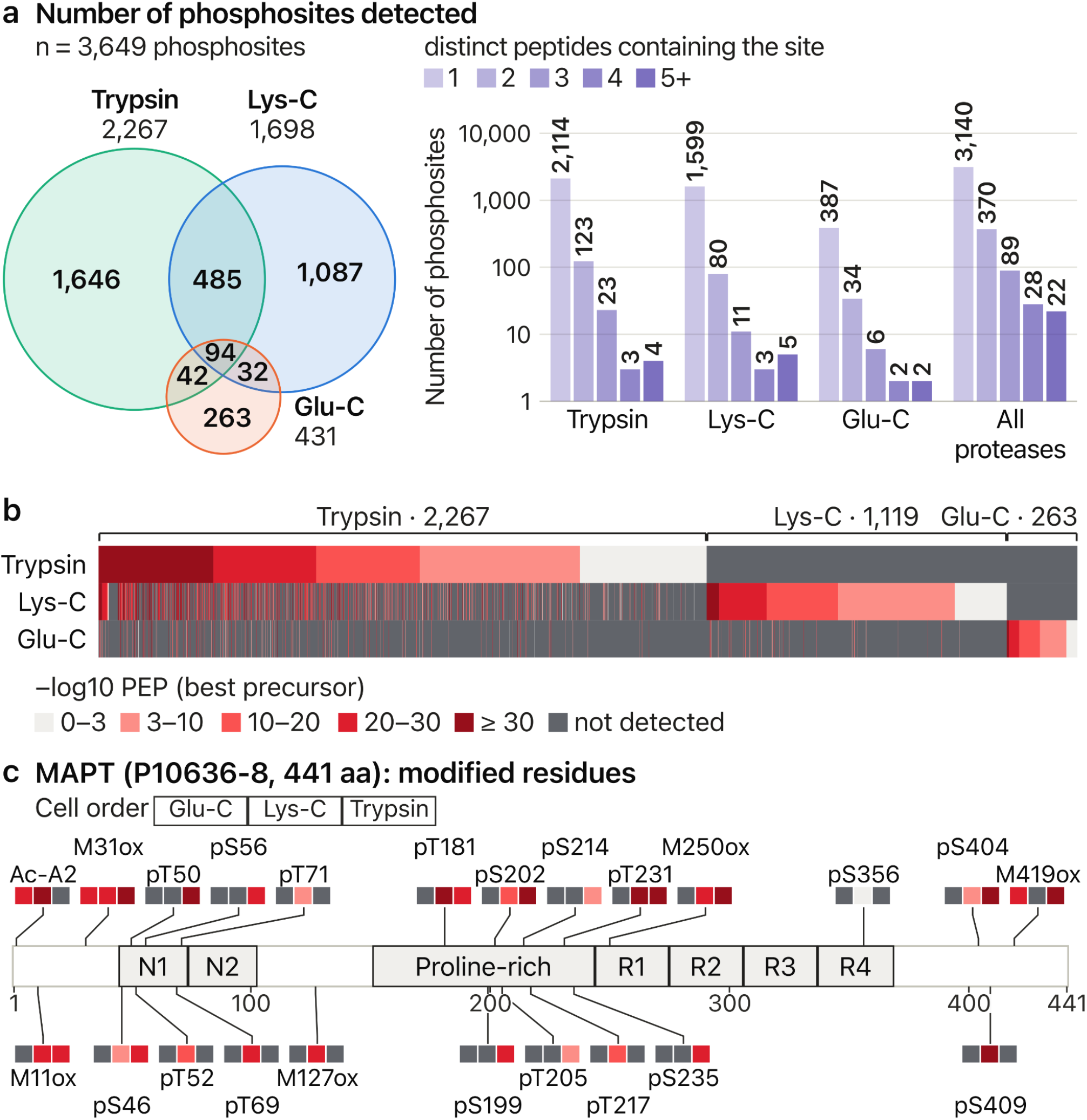
Phosphosites detected without enrichment. (a) The 3,649 phosphosites by the digests in which they were detected (left) and by the number of distinct peptides containing each site (right). **(b)** All phosphosites by protease, colored by the posterior error probability (PEP) of the best precursor; about 82% were detected by a single protease. **(c)** Modified residues of MAPT on its 2N4R 441-residue isoform (P10636-8), one cell per protease, colored as in b; N1 and N2 are the inserts encoded by exons 2 and 3, and R1–R4 the microtubule-binding repeats, R2 encoded by exon 10.

Detecting a phosphosite on multiple distinct peptides, whether within one digest or across digests, increases confidence in its identification, localization and quantification: each peptide contributes its own fragment ions, its own localization of the phosphate group and its own abundance estimate (Fig. 3a). The number of phosphosites detected on two distinct peptides rose from 123 with trypsin alone to 370 with all proteases, and the number detected on three distinct peptides rose from 23 to 89.

Detection of disease-relevant phosphosites greatly benefits from using multiple proteases. Tau has multiple phosphorylated residues^33,34^. Accordingly, multiDIA allowed mapping 17 phosphosites on its 441-residue 2N4R isoform, including the p-tau epitopes pT181, pS202 and pT231, each of which was quantified in all 10 donors with complete data by both trypsin and Lys-C (Fig. 3c). Six tau phosphosites were detected only by Lys-C, among them pT217 (in one donor) and pS409 (Fig. 3c).

## Discussion

Using the union of complementary peptide populations, multiDIA substantially increases proteome sequence coverage beyond that achievable with a single digest analyzed by DIA without extensive fractionation (Fig. 1). We previously used two proteases to increase the robustness of quantifying protein synthesis and degradation in mice^35^, and here we demonstrate the ability of multiDIA to resolve and quantify proteoforms that play central roles in neurodegenerative diseases (Figs. 2 and 3). Despite the small size of our cohort, multiDIA resolved expected decreases in dopaminergic enzymes in PD, as well as differential ratios of proteoform to canonical protein in PD versus control (Supplementary Figs. 1 and 2). We expect that larger cohorts will substantially increase the statistical power for detecting disease-associated changes in proteoform abundance, including proteolytic transactions and post-translational modifications, such as phosphorylation.

These gains in sequence coverage and identification confidence come at the cost of a 3-fold increase in instrument time per sample. This reduced throughput could be offset in future experiments that combine multiplexed DIA (plexDIA)^36–38^ with multiDIA, so that multiple samples digested with the same protease are labeled with mass tags and analyzed together. Throughput could also be increased by multiplexing samples in time using timePlex^39^.

Analyzing mixed digests from multiple proteases has been proposed as a protease-multiplexing approach for increasing coverage^40^. The idea is attractive because it exploits the ability of DIA to analyze more precursors within the same number of MS2 windows, somewhat analogous to plexDIA. However, applying protease multiplexing to three cell lines identified fewer than 45,000 peptides per sample, fewer than are identified from tryptic digests alone on older instruments, such as the Exploris 480^35,41^. In our hands, DIA analysis of mixed digests with existing computational approaches did not substantially increase the number of identified peptides. In principle, such an increase should be possible, for example with dedicated computational approaches optimized for multiplexed DIA spectra^42^.

Although multiDIA does not reach the sequence depth achieved by the most extensive multidimensional fractionation workflows, it achieves, to our knowledge, the deepest coverage attained with 3 hours of analysis time and less than a microgram of protein per sample. These characteristics make multiDIA an attractive approach for acquiring quantitative, proteoform-resolved datasets from large cohorts.

## Methods

### Sample Preparation

All postmortem frozen brain tissue samples were obtained from the Massachusetts Alzheimer’s Disease Research Center (ADRC) brain bank. A cohort of 12 donors (6 PD and 6 controls) was selected on the basis of primary and secondary diagnoses, comorbidities, and clinical and demographic characteristics. Postmortem evaluation was used to classify cases on the basis of PD and Alzheimer’s disease (AD) neuropathologic changes, including Braak staging of α-synuclein and tau pathology, Thal amyloid plaque staging, CERAD neuritic plaque staging, and the presence of TDP-43 pathology. The complete cohort demographics are shown in Table 1.

Bulk tissue processing was performed as previously described^21^. Briefly, for each case, approximately 50 mg of dorsolateral striatum was dissected and lysed using bead-based tissue disruption. Tissue lysis was performed by transferring samples to a microcentrifuge tube containing zirconium oxide beads (Next Advance 430917) and 600 μl of lysis buffer on ice. The lysis buffer contained 75 mM NaCl, 50 mM EPPS (pH 8.5), 10 mM sodium pyrophosphate, 10 mM sodium orthovanadate, 3% SDS, 10 mM PMSF, and one EDTA-free protease inhibitor tablet (Roche 11873580001). After adding the tissue, the tubes were transferred to a Mini-Beadbeater 16 (Biospec). Samples were homogenized for 30 seconds and then placed on ice for 2 minutes; this cycle was repeated three times to ensure complete tissue lysis. Samples were treated with benzonase to degrade nucleic acids, then denatured in SDS by heating. For each sample, 100 µg of protein was reduced, alkylated, cleaned up with SP3 and digested overnight at 37 °C with Lys-C, trypsin or Glu-C (enzyme:substrate ratio 1:25). Digested peptides were dried and reconstituted in 0.015% DDM and 2% formic acid for LC-MS/MS injection.

### LC-MS/MS Acquisition

Peptides were separated by nanoflow reversed-phase chromatography on a Vanquish Neo UHPLC using a 25 cm × 75 *µm* IonOpticks Aurora Series UHPLC column (AUR2-25075C18A) with a 65-minute run (55-minute active gradient) at 200 nl/min, with a standard water/acetonitrile mobile phase pair. After sample loading and a brief ramp, peptides were eluted across a shallow organic gradient, followed by a high-organic wash and column re-equilibration.

Peptides were analyzed on a timsTOF Ultra mass spectrometer operated in dia-PASEF mode. Ion mobility was scanned across a 0.75–1.3 V·s/cm^2^ range, with 50-ms ramp and accumulation times. Twenty MS/MS ramps spanning sixty precursor isolation windows were used to cover an m/z range of 350–1250.

### Data Analysis

Raw dia-PASEF data were searched directly against the UniProt human sequence database including isoforms (2024-01-01; 42,458 entries) using the directDIA workflow in Spectronaut (v21.1), without a prebuilt spectral library. The default BGS Phospho PTM Workflow was used, with the decoy generation rules modified to reflect the protease used for each digest and the quantification normalization filter type set to *None*. Standard precursor and protein false discovery rate (FDR) thresholds (1%) were used, together with the default PTM localization filter (0.75). The full Spectronaut search settings file is deposited with the MS data in MassIVE.

## Data and Code availability

Raw data, search results and settings are available at MassIVE: MSV000103483. Software, data-analysis pipelines and other supporting documentation are available at multidia.slavovlab.net and github.com/SlavovLab/multiDIA.

## Acknowledgements

We thank V. Ondruska for help with timsTOF optimizations and members of PTI and the Slavov laboratory for discussions and suggestions. We also thank D. Hornburg and J. Krieger for constructive feedback.

## Author contributions

Study design, supervision and funding acquisition: N.S., M.C., B.T.H. Sample preparation: K.A., D.O; Data acquisition: C.F. Data analysis: O.K and N.S.;. Initial draft: N.S. Writing: all authors. All authors approved the final manuscript.

---

**Supplementary Figure 1.**
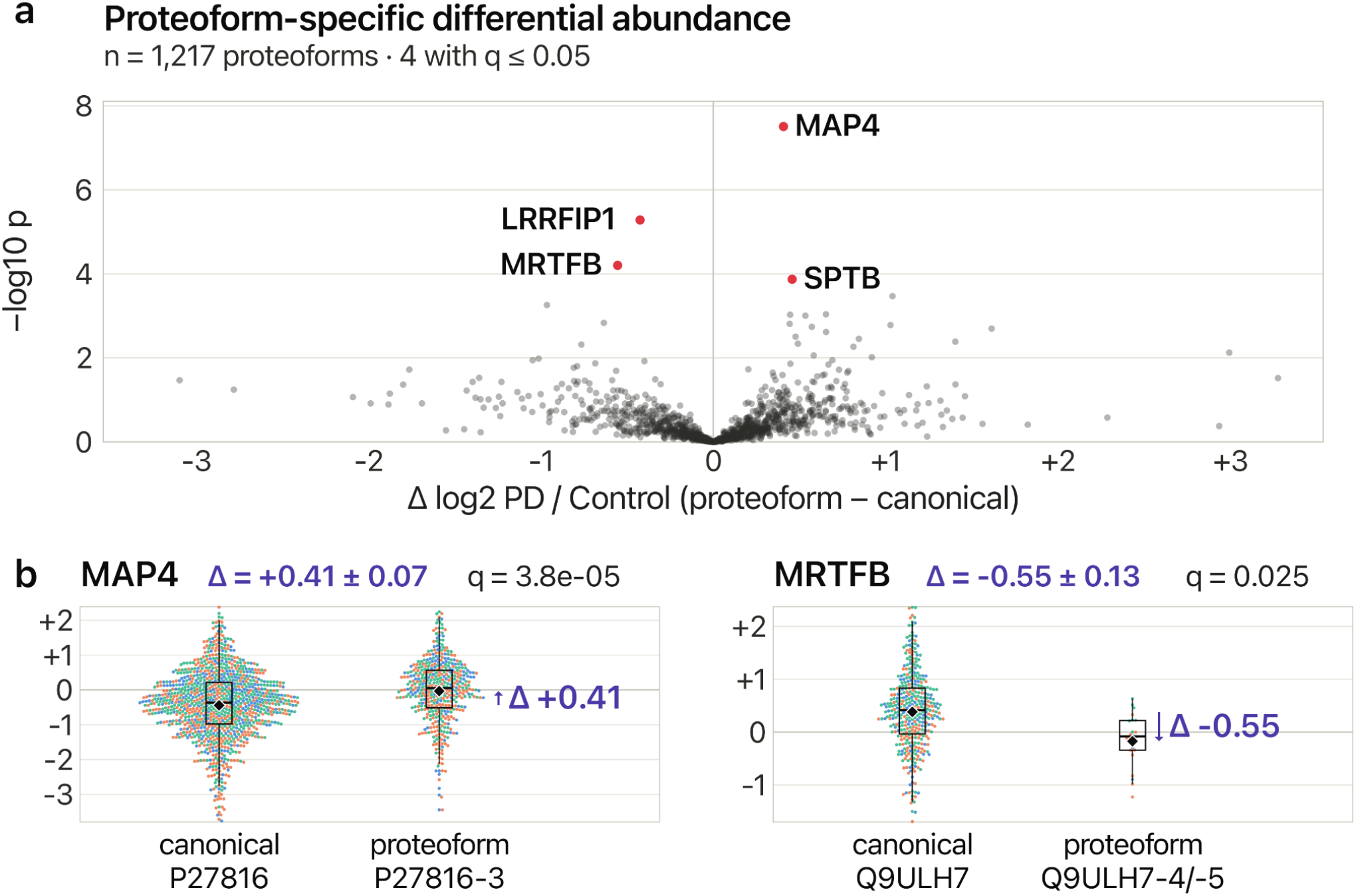
Proteoform-specific differential abundance. (a) Change in each proteoform’s abundance in LBD relative to its canonical form: Δ = (proteoform log2 LBD/control) ™ (canonical log2 LBD/control); proteoforms whose canonical form was not measured were not tested, and the four that diverge from their canonical form at Benjamini–Hochberg q ≤ 0.05 are highlighted in red. **(b)** Two of them (MAP4, MRTFB): log2 LBD/control ratio of each peptide in each LBD donor, colored by protease.

**Supplementary Figure 2.**
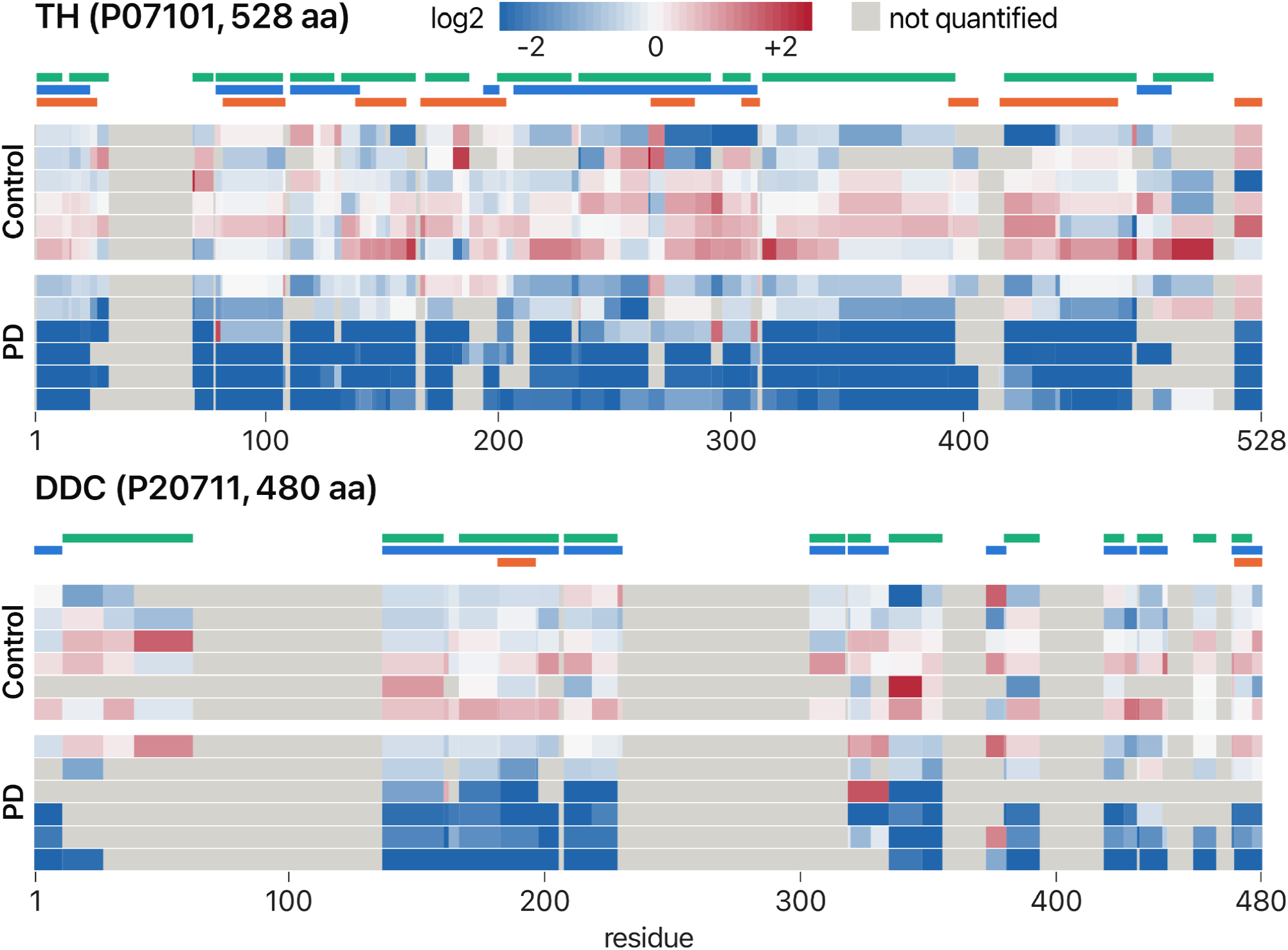
Dopaminergic enzyme peptide abundance by donor. Tyrosine hydroxylase (TH) and DDC peptide abundance along each protein, as in the top of Fig. 2c: one row per donor, ordered within each group by profile similarity; each residue shows the mean log2 abundance of the peptides covering it, relative to the control mean (each control against the other controls); grey, not quantified. Bars above the rows mark trypsin (green), Lys-C (blue) and Glu-C (orange) peptides.

**Supplementary Figure 3.**
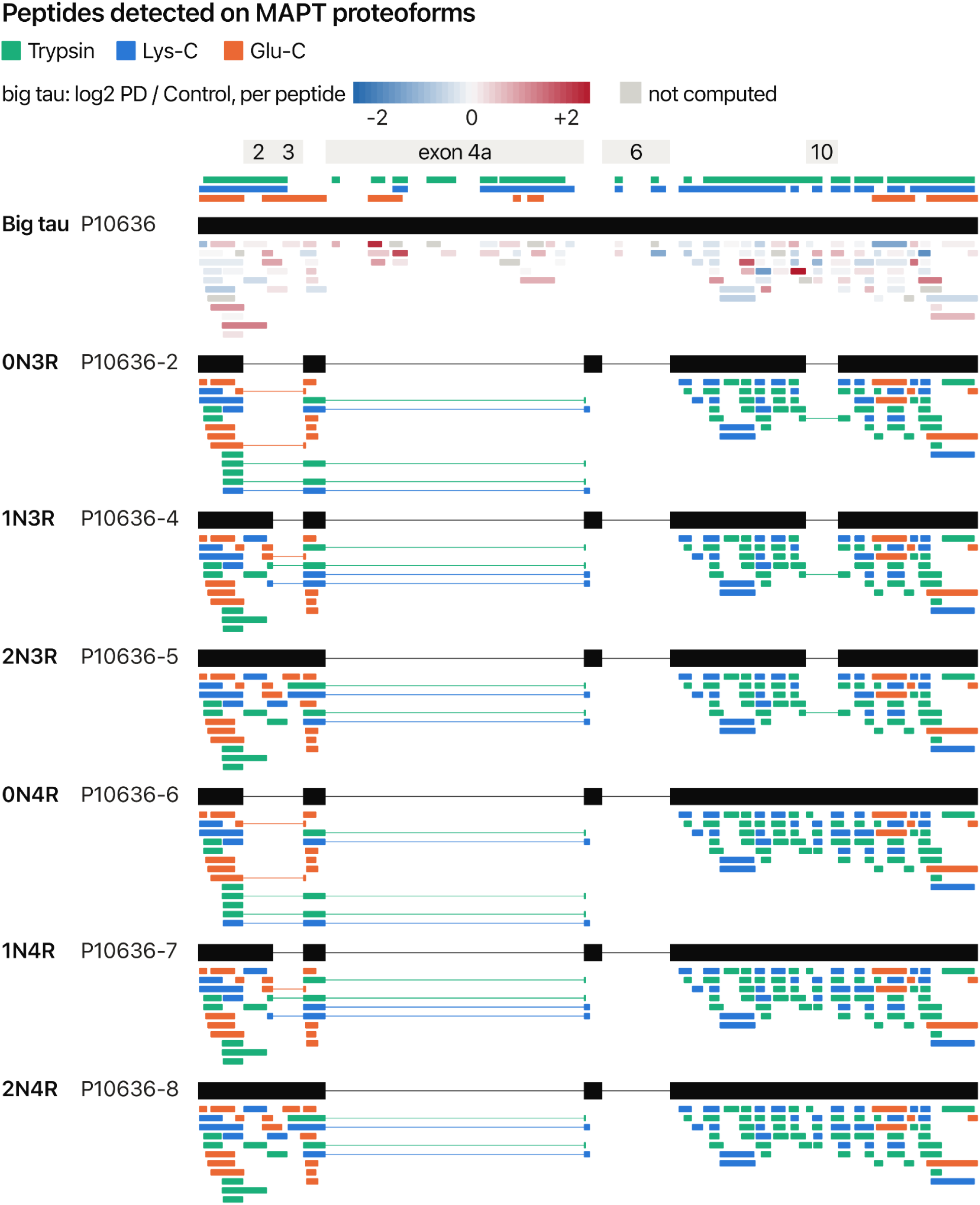
Peptides detected on MAPT proteoforms. Big tau (P10636) and the six adult brain proteoforms (0N3R to 2N4R, P10636-2 to -8), aligned on the big-tau sequence: black, residues each proteoform contains; thin line, exons it skips (exons 2, 3, 4a, 6 and 10 marked above). Bars above big tau mark the residues covered by each protease’s tau-unique peptides. Below each proteoform, the peptides unique to tau that occur in its sequence, colored by protease; a peptide drawn across a gap spans the junction of the skipped exon. Big tau’s peptides are colored instead by log2 PD / control, each peptide against its own control mean as in Fig. 2c; grey, not computed.

